# Recoveries of Ringed Red-Footed Falcons (*Falco vespertinus*) in Germany - An update documentation

**DOI:** 10.1101/2020.09.11.293910

**Authors:** Dietmar Kunze, Lisa Nennstiel

## Abstract

Bird ringing is used for a long time for scientific investigation of migration routes and to better understand breeding events as well as population ecological aspects. It is applied as an inexpensive method although the well-known and major disadvantage of bird ringing is the usually low response rate. In the case of rarer species such as the Red-Footed Falcon (RFF), however, this quota could be higher due to the exclusivity and greater attention of observers. Motivated by own field observations of color-ringed RFF south of Brunswick, Germany in 2019 and no clear and comprehensive publication of verified additional recoveries and ringings, we did further research regarding this issue by following methods: 1) Contacting European Bird Ringing Centers and associated projects, 2) Query and comparison of files with the three national German Bird Ringing Stations, 3) Expanded literature research, 4) Evaluating reports in the citizen science platform ornitho.de, 5) Checking websites of ringing projects (in particular satellite tracking programs) and 6) Own observations.

Surprisingly, this study revealed 18 recovered RFF ringed in foreign countries (14 Hungary, 3 Italy, 1 Romania (GPS tracked bird)) during migration or post-nuptial pre-migration time. Additionally, 1 RFF that was caught and ringed in Germany was recovered abroad.

This result updates and increases the number of recoveries of RFF in Germany compared to the actual published state on the order of 18 (so far none) and on the order of 6 concerning ID-encoded RFF (GPS-bird excluded) compared to documentation state of the three Bird Ringing Centers in Germany (so far 3 at Beringungszentrale [BZ] Hiddensee, 1 at Institut für Vogelforschung [IfV] Heligoland). Our research and evaluation of raw data succeeded to a 100% identification rate of the bird’s origin countries (n=18) while the rate of ID-encoded RFF by color ring codes revealed 58.8% (n=10, GPS-bird excluded). The reported-by-observer response rate was 41.2%. Interesting data of about the phenology, age and origin of the RFF recovered in Germany are presented. Questions and considerations about the recent reporting system of ringed birds and the increased numbers of RFF during the last years in Germany are discussed.

## Introduction

The last two decades, Red-Footed Falcons (RFF, *Falco vespertinus*) appeared every year in Germany either in spring during their loop migration back from Africa or in their post-nuptial pre-migration period as citizen science data (ornitho.de), regional annual reports and data of rarity commissions reveal. Even the decades before, RFF were probably quite regular visitors in Germany, but data situation does not allow a reliable comparison due to methodological aspects. In addition, when evaluating older sources, special attention must be paid to the aspect of confusing 1cy RFF with the Eurasian Hobby (*Falco subbuteo*), so it is not unrealistic that a certain number of RFF were overlooked at that time (e.g. von Transehe & Schüz 1940). In some years, numbers of RFF in Germany increase to invasive-like magnitudes, as happened in e.g. 1927 (Niethammer 1938), 1968 (Heldt 1969), 2008 and 2015 (König et al. 2015). Also 2016 and 2019 were years with high numbers referring to evaluated data of ornitho.de. Regarding this subject, it is absolutely necessary to make a clear phenological distinction between the abundances of RFF during spring and autumn: While spring occurrences are most likely to be affected by strong NE trade winds in Africa (e.g. Palatitz et al. 2018), the late-summer/autumn phenomenon in the post-nuptial period is still not fully understood.

Recoveries of two ringed RFF in late Summer 2019 S of Brunswick in the N foreland of the Harz mountains and no entries of recoveries in the current Atlas of Bird Migration (Bairlein et al. 2014), have been the motivation of this actual research in attempt to gather all available data of ringed RFF that have been seen in Germany the last decades to obtain a clear and comprehensive documentation.

## Methods

Data were obtained through the following methods: 1) Query and comparison of files with European bird ringing stations or their partner organizations, respectively, which hold breeding populations of Red-Footed Falcons and/or conduct ringing schemes for this species: Contacted countries were Austria, Belarus, Bulgaria, Estonia, Finland, France, Hungary, Italy, Latvia, Lithuania, Poland, Romania, Russia, Serbia, Slovakia, Spain, Switzerland Portugal and Ukraine, 2) Query and comparison of files with the three national German Bird Ringing Stations Hiddensee, Radolfzell and Heligoland, 3) Expanded literature research, 4) Review of recovery reports in the citizen science project ornitho.de, 5) Checking websites of ringing projects (in particular satellite tracking programs), 6) Own field observations. Data research ended on 14/06/2020.

Referring 4), the ring codes were either already encoded by the observers but not yet reported to the responsible national ringing centers or were attempted to encode by the first author with support of the respective ringing projects on the basis of observers` pictures. Regarding RFF ringed in Germany we followed Bairlein et al. (2014) defining recoveries for migratory species >100 km.

## Results

### Recoveries

A total of 18 in Germany recovered RFF were found in this research (Tab.1, Fig. 1). This result updates and increases the number of recoveries of RFF in Germany compared to the actual published state on the order of 18 (so far none) and on the order of 14 from the documentation state of the three Bird Ringing Centers in Germany (so far 3 at BZ Hiddensee, 1 at IfV-Heligoland/Wilhelmshaven).

**Tab. 1:**
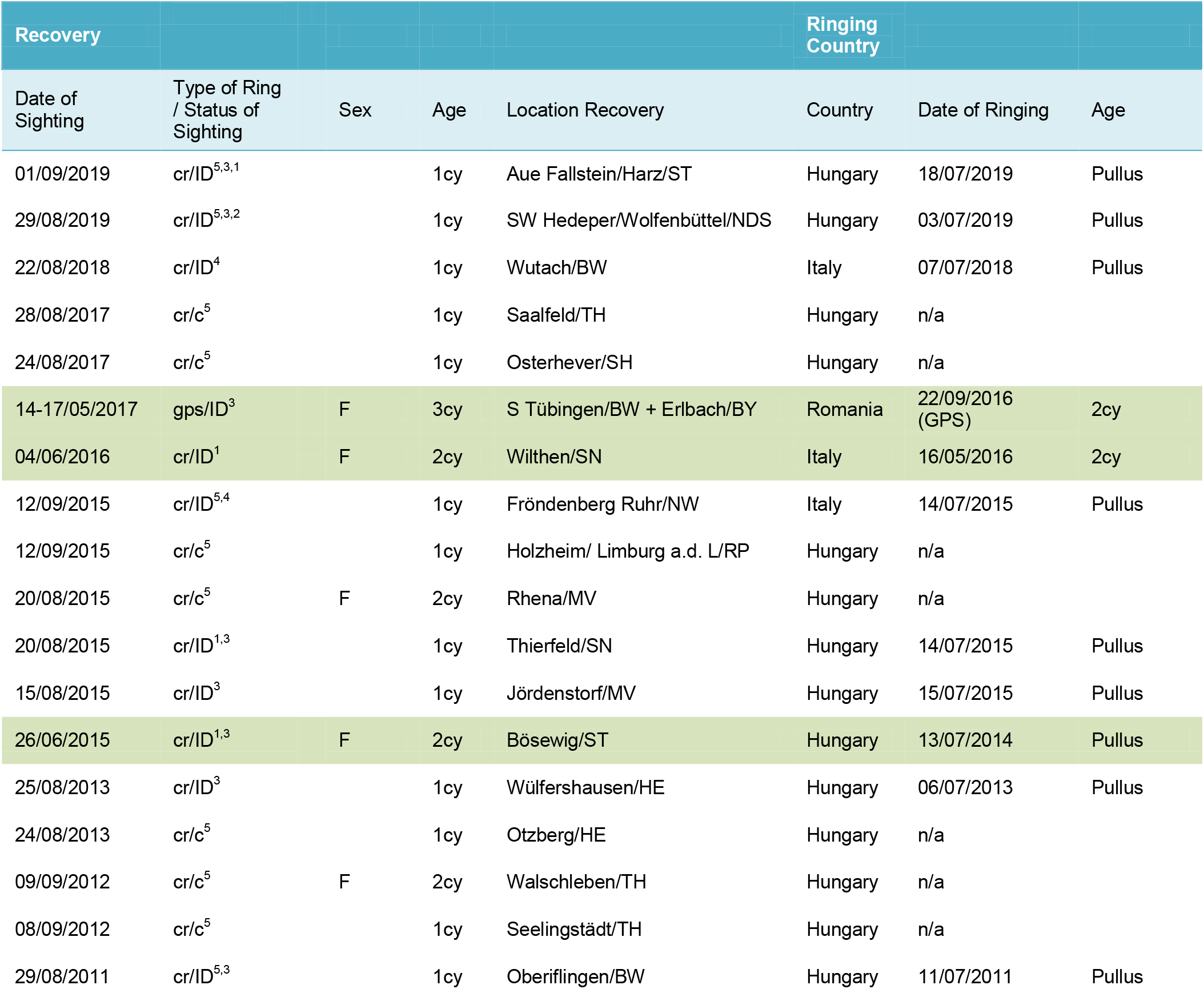
All known and evaluated recoveries of Red-Footed Falcons (*Falco vespertinus*) in Germany until of 14th June 2020. Birds with ID (Ring number encoded) are shown with day of ringing, country and age of ringing. For region see map (Fig.1). Type of ring/Status of sighting means: cr = color ring, c = only country known due to color ring scheme and positions. Pale green highlighted: Spring recoveries. Data source: ^1^ Beringungszentrale Hiddensee, ^2^IfV Helgoland/Wilhelmshaven, ^3^MME Hungary, ^4^Lipu-BirdLife Italia, ^5^evaluated data from ornitho.de (sum n=18).

**Fig. 1:**
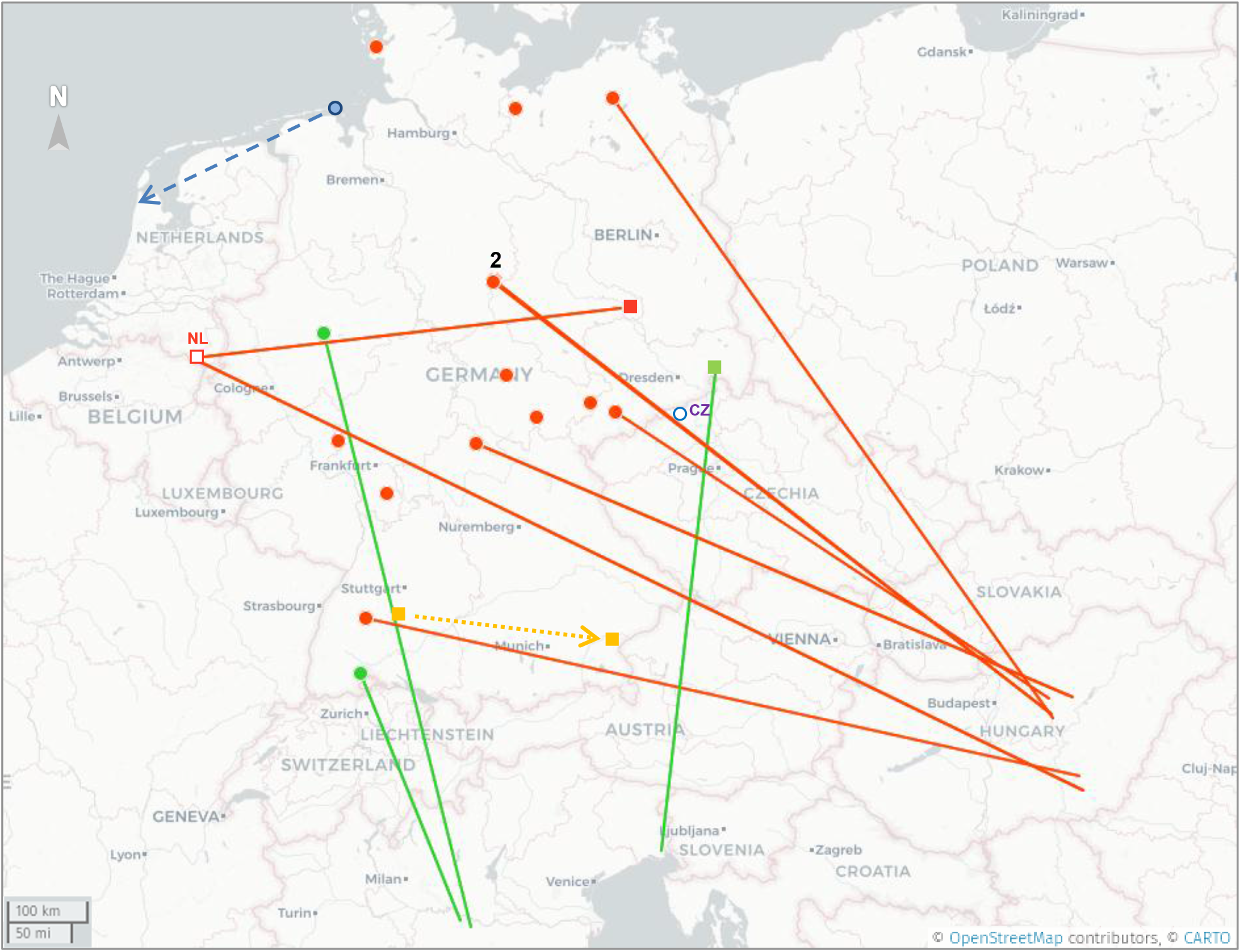
All known and evaluated recoveries of Red-Footed Falcons (*Falco vespertinus*) in Germany until 14^th^ of June 2020 (n=18). Solid lines depict recoveries with exact origins of ringing: Line end without a dot or square depicts the place of ringing (all places of breeding population except the 2cy Italian bird near border to Slovenia), line end with a dot or square shows the place of recovery (n=10). Without lines: Only the country of origin is known (n=7). Dots: Late summer/autumn post-nuptial pre-migration (2 over dot means two different birds nearby) (n=15). Squares: Spring migration (n=3). Colors signify the country of ringing/breeding population; Orange: Hungary (orange framed white square is a pre-recovery in the Netherlands (NL)), Green: Italy, Yellow: Romania (GPS-bird, dotted yellow arrow between two spring stopovers in Germany), Blue: Germany (only one ringing with recovery abroad, namely The Netherlands in late summer 1938; dashed blue line to place of recovery: Callantsoog, North Holland), White blue framed dot (CZ): Recovery of a RFF in the todays Czech Republic only a few kilometers away from the German border, ringed as 2cy in spring 1965 (Tunisia) and recovered in October 1977 (CRBPO 2019 *in litt.*). Map: Based on ©OpenStreetMap contributors, ©CARTO.

In addition, only one case of a RFF ringed in Germany and recovered >100 km away was found in literature: A 1cy bird ringed on 08/08/1938 on the German barrier island Wangerooge, today Lower Saxony, observed again 10 days later and 235 km WSW at Callantsoog, North Holland, Netherlands (cf. Großkopf 1968).

Extensive literature evaluation and comparison of the data with the French Bird Ringing Center (CRBPO) revealed an interesting recovery in the todays Czech Republic only a few kilometers away from the German border (Fig. 1). This RFF is most likely the bird assigned to Germany in a recent publication probably due to the small scaled not very detailed map in the original reference Haraszthy (1982) (cf. Bird Study, 63(3), 406-412) and was ringed of age +1 year during spring loop migration at El Haouaria, Nabeul/Cab Bon, Tunisia on 20/05/1965 by French ornithologists, recovered 12 years later in Záluží (Litvínov) N of Brix, former Czechoslovakia, present Czech Republic on 23/10/1977 (sic!) (CRBPO 2019 *in litt*, Fig.1). This female bird is of an unknown breeding population and documented with >13.25 years as the age record in the EURING longevity list (EURING 2017).

### Origins and Encoding

The vast majority of recoveries were ringed in Hungary (14), followed by Italy (3) and Romania (1, GPS track evaluation, Tab.1). All of the latter birds were color ringed and from breeding populations of their origin countries. 17 out of the 18 recoveries were made by field recognition, description and/or pictures of these color rings. One bird was satellite-tracked and was only detected by its GPS location protocol crossing S Germany with stopovers from 14-15/05/2017 and 15-17/05/2017 in a far west loop during spring migration on the way to its 2017 breeding ground SW of Odessa, Ukraine (cf. satellitetracking.eu, falcoproject.eu, Fig.1, Tab.1). The GPS transmission took place during the EU LIFE+ project “Conservation of the Red-footed Falcon in the Carpathian Basin (LIFE11/NAT/HU/000926)” (MME 2012-2018). The only recovery of a solely metal-ringed bird remarks at the same time the only remote finding of one of the very few RFF ringed in Germany: The Wangerooge bird of 1938.

While 10 birds could be safely encoded (GPS-bird not included due to methods; 7 Hungary, 3 Italy), another 7 ringed RFF could not offer secure details to determine individual status (Fig. 1).The rate of encoded RFF rings is therefore 58.8%. Hence, 41.2% were not encoded or could not be encoded safely in review of data, respectively. The ratio of encoded/not-encoded rings of RFF results 1.43. The reported-by-observer response rate was 41.2% compared to all observations of RFF that were documented in the citizen science platform ornitho.de and the lists of all German and European bird ringing stations mentioned before.

### Phenology

15 out of 18 RFF were reported during late-summer/autumn in the post-nuptial pre-migration time, 2 RFF were recovered during spring migration (1 Italian, 1 Hungarian) supplemented by the Romanian RFF revealed by satellite tracking.

The year with the most recoveries of ringed RFF was 2015 (n=6), followed by 3 documentations in 2017 (incl. GPS-bird) and 2 recoveries each for 2012, 2013 and 2019. Data of 2011, 2016 and 2018 exhibited 1 recovery each (Fig. 4, Tab.1).

Recoveries exhibited 13 birds of the first calendar year (1cy), 4 of the second calendar year (2cy) and 1 of age 3cy (Fig. 3). In the post-nuptial pre-migration time only 2 birds were of age 2cy while 13 were of age 1cy. During spring migration, 2 birds of age 2cy and 1 bird of age 3cy (GPS) were determined (Fig. 2, Tab.1). Regarding the sex of the recovered RFF, only the birds older than 1cy could be of safe determination. All 5 birds of age >1cy were female.

**Fig. 2:**
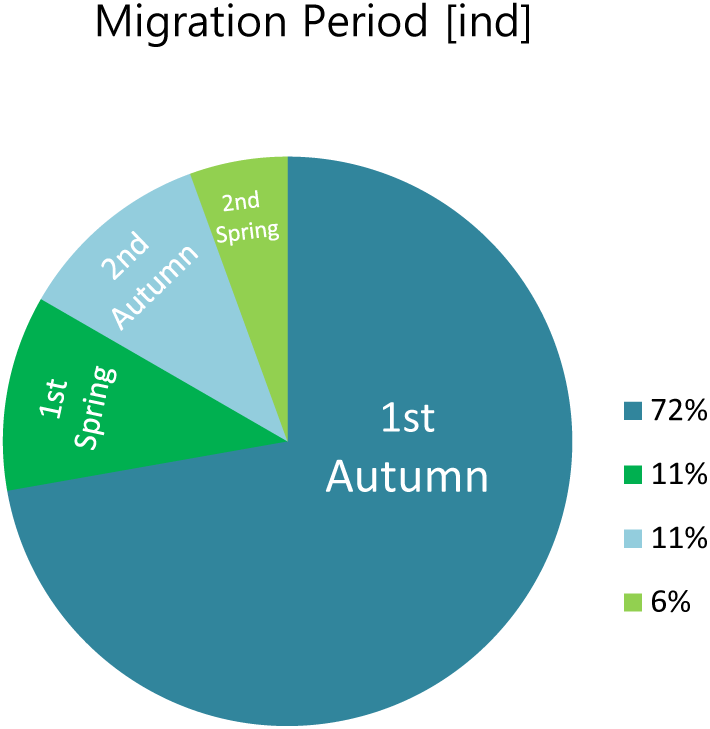
Share of rediscovered ringed Red-Footed Falcons (*Falco vespertinus*) on individual basis in the respective migration period in Germany until of 2019 (n =18).

**Fig. 3:**
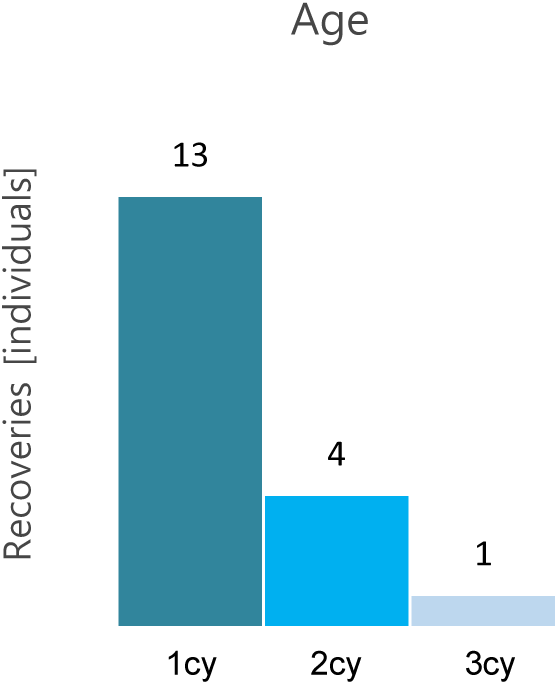
Share of calendar year age (cy) of all recovered individuals in Germany (n=18) until of 14/06/2020.

All birds treated here were not yet adult at the time of ringing (n=14, 77.8% as pullus or 1cy; n=2, 11.1% ringed as 2cy). The recoveries also took place exclusively immediately in the ringing year or in the following year. Concerning the age of RFF sighted at the migration periods, data exhibited that this was 72.2% during the birds’ first pre-migration (post-nuptial period, n=13, 1cy) and 11.1% during the first spring migration (n=2, 2cy), while during the birds’ second autumn pre-migration it revealed 11.1% (n=2, 2cy) and regarding the second spring migration this was 5.6% (n=1, 3cy GPS bird) (sum n=18, Fig. 2). None was seen in consecutive years (as adult). Likewise, none of the ringed birds were read at more than one location in Germany, except the two solely technical registered stopovers of the GPS bird.

## Discussion

### Ringing

Bird Ringing is used for over 120 years now for the long way to understand migration related behavior and population ecological aspects as well (e.g. Bairlein 2001). Nowadays, along with technical progress very precise GPS-based modules enabled the application even for smaller birds for migration monitoring (e.g. Fiedler 2009). Technical equipment is getting more and more tiny and effective due to new knowledge in the semiconductor sciences. Pedersen et al. (2020) recently even satellite tracked and studied the long-distance migration of a songbird namely the Red-backed Shrike (*Lanius collurio*) with this update technology. However, the (color) ringing is currently still applied as a very inexpensive method although the well-known and major disadvantage of bird banding is the usually very low response rate. Hence, a present and realistic compromise that describes a low-cost, bird convenient and also effective in terms of feedback rate method is desirable. The gap between the time when the satellite modules will be ready to use in ring size and still be affordable and the status quo can therefore best be bridged by optimizing the reported-by-observer feedback rate of color rings.

Even if probably most of the recoveries of ringed (song) birds are made by the ringers themselves through targeted catching at their projects’ sites, random large-area covered observations by birdwatchers are very important and also welcomed by the ringers. The latter item especially concerns larger species like the RFF when rings are relatively easy to read or color ringing patterns reveal the place of ringing. Although sightings are supposed to be reported to the corresponding national Bird Ringing Centers namely the *Institut für Vogelforschung (IfV) - Vogelwarte Helgoland*, the *Beringungszentrale Hiddensee* and the *Max-Planck-Institut für Verhaltensbiologie* (former *Vogelwarte Radolfzell*), this procedure is probably not yet known by all observers.

As the number of amateur and spare time ornithologists fortunately rises and because most birders in 2020 use the very convenient and one-address centralized portal ornitho.de, it would be very effective to provide a filter or even contact form system in this most-used bird report database in Germany. In the authors’ opinion, this portal-based way would be a great benefit to the ringing centers and all ringing projects in Europe. As this study shows, the response rate of color ringed birds like the RFF could increase by almost 60%. Citizen science observers’ data would, in the best case, go directly to the according ringing station and project as well. It also would prevent data from being lost and, in busy times like that, many mails and research work would be redundant. Hence, this study in hands suggests a necessary action to improve the current system of recovery reports concerning recoveries of all ringed bird species by the latter proposals.

### Classification of results

Since this is a documentation of ring recoveries of RFF in Germany, the quantity of data is naturally not sufficient enough to exhibit reliable interpretation or even statistical work. Nevertheless, the records should be regarded in context with the current knowledge of migration and dispersion of this interesting species.

### Spring migration

The vast of spring migrating RFF probably pass the Mediterranean Sea via mainland Italy (synchronized migration counts, Premuda et al. 2005) and/or neighboring islands (satellite tagged RFF, MME 2012-2018). Albeit, the latter GPS program also revealed routes via the Greek and Balearic islands (scarce migrant of Mallorca, Rebassa et al. 2013).

Spring abundances in Germany are presumably most influenced by strong trade winds and other aggravating factors in Africa during the RFF’s exhausting loop-migration that drift away the birds in a too far NW direction (c.f. Palatitz et al. 2018). That this detour in migration more often concerns young birds is not entirely proven yet (c.f. Solt 2018). However, the German springtime recoveries of ringed RFF could support the latter thesis (Fig. 2, 4; Tab. 1).

**Fig. 4:**
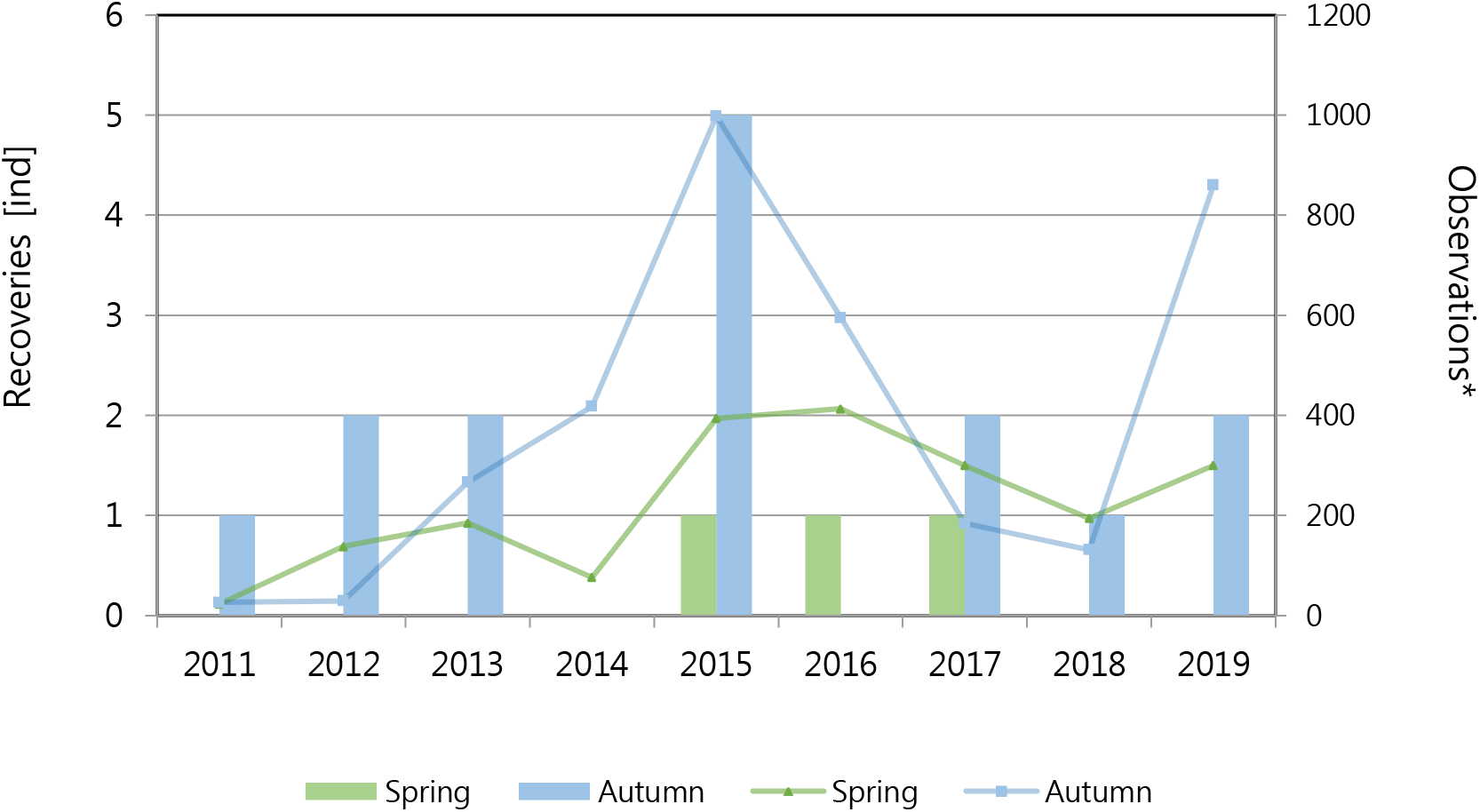
Recoveries of RFF in Germany on individual status in spring (green columns) and autumn (blue columns) of the years 2011-2019 (n=18) in comparison to numbers of observations* in Germany generated in ornitho.de (green line: spring, blue line: autumn). *Note: Numbers do not depict individuals! Numbers are generated in the citizen science platform ornitho.de and include repeated and multiple sightings. Data are not reviewed systematically yet.

### Post-nuptial dispersal

Long-time studies and satellite tagging of RFF revealed that RFF break up family bindings usually within 3-4 weeks after fledging, sometimes even before the migration period begins (Palatitz & Solt 2018). In this pre-migration dispersal phase, young birds are highly mobile and occasionally head to seemingly unfavorable N or NW directions several hundreds or even over 1000 km from their breeding colonies (e.g. Palatitz & Solt 2018, Palatitz et al. 2018). A similar phenomenon, albeit probably not of this size, is also known from the Lesser Kestrel (*Falco naumanni*): in Bounas et al (2018) these birds flew long distances (>500km) to their roost sites during their post-nuptial pre-migration time, partly also northwards. Both species also gather to large groups at their roost sites after the breeding period.

In some years, particularly high numbers of RFF can be observed in Germany (also in Poland, cf. Golawski et al. 2017). These irruptions are still not fully understood. However, weather conditions and aspects of food supply are discussed (e.g. Golawski et al. 2017, Palatitz & Solt 2018). Our presented recoveries of ringed RFF in Germany support previous observations and earlier reports (e.g. Heldt 1969) that increased abundances of 1cy RFF (less 2cy and adult) occur in the post-nuptial pre-migration period in Germany.

In addition to the above described hypothesizes, an explanation or at least a connection with a good breeding success of the RFF and its subsequent dispersion would be obvious. However, regarding the data of the large RFF population in Hungary and Romania, at least the sampled monitoring areas of the LIFE+ projects, show that 2015 was a year of extraordinary poor breeding success with particularly low offspring (Palatitz et al. 2015, 2018). The most likely cause was, besides bad weather (high precipitation), the total collapse of the common vole (*Microtus arvalis*) population after an extreme gradation the previous year (Palatitz et al. 2014, 2015). Hence, the influx of RFF in Germany in 2015 could be driven by a particularly strong food pressure and the evoked social population ecological aspects of this gregarious species with rather complex social behavior (cf. Palatitz et al. 2018). Special wind extremes for 2015 in Hungary that could support a thesis of the birds’ NW drifting are at least not known for this post-nuptial phase (www.timeanddate.com, www.worldweatheronline.com). However, prevailing wind direction and also less striking characteristics in wind speed or weather conditions could play a role anyway (cf. Golawski et al. 2017).

The occurrences of 2cy RFF could be explained by the fact that the latter usually do not yet form pairs or breed (e.g. Solt 2018a,b) and therefore accompanied the 1cy birds in their dispersion. Regarding the rarer observations of adult RFF during that pre-migration phase, it might be possible that those birds had no breeding or pair building success that year and hence wandered around longer distances: Solt (2018a,b) describes that female RFFs exhibiting stronger pronounced head and body patterns were dominant against paler and buff plumage females and also that late breeding ground arrivals had worse chances to breed successfully. However, more studies need to be done about that issue to get reliable explanation.

During the autumn 2019 influx, when the two RFF ring records of the same breeding population in Hungary (Jaszbereny, Jasz-Nagykun-Szolnok) were observed in our region S of Braunschweig/Brunswick (Tab.1, Fig.1) while a year with a strong gradation of the common vole (*Microtus arvalis)* occurred. Thus, the falcons faced a very good food supply and could be observed feeding on lots of voles several times. The weather was also fine during the documented time from late August (day: 32-33° C, night: 17-23 °C lowest, low windspeeds 12 km/h max, changing direction, mostly S and N, no precipitation) and the first week of September before commencing their migration (day: 17-24° C max, night: 8-14°C lowest, windspeed 4-26 km/h max, all directions except E, no precipitation) (www.timeanddate.com). In Hungary, after the break-down of the common vole population in 2015, numbers of the latter seemed to have recovered in 2017, at least in the northern breeding colonies of the RFF (cf. Palatitz et al. 2017, Somogyi & Horváth 2019). The authors of this paper in hands are not aware of any more current figures concerning the Hungarian population of *Microtus arvalis* as also any data concerning the breeding success of the RFF for the year of 2019.

Overall, the increased abundances of RFF in Germany have been striking in recent years and are also described with high numbers in Poland (Golawski et al. 2017). As also Golawski et al. (2017) hypothesized, stochastic reasons probably plays a role here: The important RFF populations in Hungary and West Romania have grown again strongly after the most threatening decline in earlier years. In Hungary alone starting from 2006 of only about 600-700 breeding pairs (BP) numbers rose to the actual range of 1150-1350 BP (Palatitz et al. 2015, 2020). This is probably largely due to the installation of several thousand artificial nest boxes and additional conservation measures in land use management during the LIFE+ projects (LIFE11/NAT/HU/000926, LIFE05/NAT/HU/122) (e.g. Kotymán et al. 2015; Palatitz et al. 2009ab, 2015b, 2018).

The cheering success of the conservation projects also is most likely the reason for higher numbers of recoveries of color ringed RFF the last decade in Germany (Fig. 4), since Hungary rings several hundreds of RFF per year: In 2018 no fewer than 702 (sic!) juveniles had been color ringed with quite a high number of 1013 incubating pairs out of 1150–1250 BP (of 1114 known territories) (cf. Palatitz 2020).

Regarding the Italian RFF, namely the two 1cy birds recovered in autumn 2015 and 2018 in Germany, both had a the rather short way across the Alps in the post-nuptial dispersion phase, assuming that they chose this route more or less directly. This occurrence is probably due to the dispersion of the northern Italian population. This artificial nest-box population in the Parma Region of about 80-85 BP now is stable for some years and was evaluated since establishing. Results are presented in Calabrese et al. (in press.).

Also this spring 2020 exhibited increased numbers of RFF in Germany regarding data in ornitho.de. Palatitz et al. 2018 presume an approx. 5-year cycle of particularly strong trade winds in Africa which support the far western drift. Further investigation of these irruptive years in Germany have to be studied by first means of observing and documenting the consecutive years. Here, the bird’s age, their diet, length of stay, weather conditions and of course possible ringed individuals need to be taken into special account to be able to carry out long-term analyzes with sufficient numbers of reliable data.

## Acknowledgements

A lot of people contributed to this documentation and we would like to thank all of them. All the contacted bird ringing centers in Europe are acknowledged for a short-term reply of the queries. Particularly we are grateful to Wolfgang Fiedler (MPI Radolfzell), Olaf Geiter (ifv Heligoland) and Christof Herrmann (BZ Hiddensee) for their friendly help referring data research. Many thanks to Attila Nagy (MILVUS Romania), Marco Gustin (Lipu – Bird Life Italy), Peter Palatitz, Zsolt Karcza (MME Hungary), Romain Provost (CRBPO) and Christoph Zöckler for their unremittingly support at assigning and encoding the recoveries. Elmar Fuchs, thank you so much for translating Russian literature. A special thanks is dedicated to Ralf Isensee who first discovered the RFF in our region in 2019. Last but not least, we would like to thank the editors of this AVES issue.

## References

Bairlein F., Dierschke J., Dierschke V., Salewski V., Geiter O., Hüppop K., Köppen U., Fiedler W. (2014). Atlas des Vogelzugs. Ringfunde deutscher Brut-und Gastvögel.

Bairlein, F. (2001). Results of bird ringing in the study of migration routes and behaviour. Ardea, 89(1), 7–19.

Bounas, A., Tsaparis, D., Gustin, M., Mikulic, K., Sarà, M., Kotoulas, G., & Sotiropoulos, K. (2018). Using genetic markers to unravel the origin of birds converging towards pre-migratory sites. Scientific reports 8(1), 1–9.

Calabrese L., Mucciolo A., Zanichelli A. & Gustin M. in press. Trend effects of nest boxes on the most important population of red-footed falcon *Falco vespertinus* in Italy. Conservation Evidence.

EURING (2017). https://euring.org/files/documents/EURING_longevity_list_20170405.pdf (last access: 10/07/2020).

Fiedler, W. (2009). New technologies for monitoring bird migration and behavior. Ringing & Migration 24(3), 175–179.

Glutz von Blotzheim, U. N., Bauer, K. M., & Bezzel, E. (2001). Handbuch der Vögel Mitteleuropas. Digital Issue.

Golawski, A., Kupryjanowicz, J., Szczypinski, P., Dombrowski, A., Mroz, E., Twardowski, M., Kielan S., Antczak, K., Pagorski, P. & Murawski, M. (2017). Autumn irruptions of red-footed falcons Falco vespertinus in east-central Poland. Polish Journal of Ecology, 65(3), 423–433.

Haraszthy, L. 1982. Kobchik. In Il’ichev (ed.). Migration of Birds of Eastern Europe and North Asia. Falconiformes andGuiformes, 164–165. Moscow Science Publishing House‘nauka’, Moscow (in Russian).

Heldt R. (1969). Zum Einflug des Rotfußfalken, *Falco vespertinus*, 1968. Corax (19), 1–7.

König, C., S. Stübing & J. Wahl (2015): Frühjahr 2015: Zugvögel im Plan, Zwergmöwen vom Winde verweht und Rotfußfalken auf Abwegen. Der Falke 2015 (8), 28–32.

Kotymán, L., Solt, S., Horváth, É., Palatitz, P., & Fehérvári, P. (2015). Demography, breeding success and effects of nest type in artificial colonies of Red-footed Falcons and allies. Ornis Hungarica 23(1), 1–21.

MME (2012-2018). http://falcoproject.eu/en/content/life (last access: 10/07/2020).

Nagy, A. (2018). Conservation of the Red-footed falcon in Western Romania. In: Palatitz, P., Solt, Sz., Fehérvári P. (Eds.) (2018). The Blue Vesper. Ecology and Conservation of the Red-Footed Falcon, Budapest, MME, 181–190.

Noga, M., Vadel, L., & Slobodník, R. (2017). Review and summary of red-footed falcon (Falco vespertinus) observations during migration periods in Slovakia. Raptor Journal 11(1), 51–67.

Palatitz P. (2020). Results of the Red-Footed Falcon (*Falco vespertinus*) population surveys in Hungary in 2018. HELIACA (16), 27.

Palatitz, P. (2018). Post-nuptial migration. In: Palatitz, P., Solt, Sz., Fehérvári P. (Eds.) (2018). The Blue Vesper. Ecology and Conservation of the Red-Footed Falcon, Budapest, MME, 117–126.

Palatitz, P. & Solt, Sz. (2018). Dispersion, independence, moult and post-nuptial roosting. In: Palatitz, P., Solt, Sz., Fehérvári P. (Eds.) (2018). The Blue Vesper. Ecology and Conservation of the Red-Footed Falcon, Budapest, MME, 103–116.

Palatitz P., Solt Sz., Horváth É., Fehérvári P., Kotymán L., Sümegi Z., Piross Imre S., Borbáth P. & Juhász T. (2017). A Kékvércse-védelmi Program éves jelentése - 2017. HELIACA 2017, 45–48.

Palatitz P., Solt Sz., Fehérvári P., Kotymán L. & Horváth É. (2015a). Kékvércse-védelmi Program évesjelentés – 2015. HELIACA 2015, 42–50.

Palatitz, P., Fehérvári, P., Solt, S., & Horváth, É. (2015b). Breeding population trends and pre-migration roost site survey of the Red-footed Falcon in Hungary. Ornis Hungarica 23(1), 77–93.

Palatitz P., Solt Sz., Horváth É., Fehérvári P., Kotymán L., Piross Imre S. (2014). A Kékvércse-védelmi munkacsoport 2014. évi beszámoló. HELIACA 2014, 12–17.

Palatitz, P., Fehérvári, P., Solt, S., & Barov, B. (2009a). European Species Action Plan for the red-footed falcon *Falco vespertinus* Linnaeus, 1766. https://ec.europa.eu/environment/nature/conservation/wildbirds/action_plans/docs/falco_vespertinus.pdf (last access: 10/07/2020).

Palatitz P., Solt Sz., Fehérvári P., Ezer Á. & Bánfi P. (2009b). Az MME Kékvércse-védelmi Munkacsoport beszámolója – a LIFE projekt (2006-2009) fôbb eredményei. HELIACA 2009, 14–23.

Pedersen, L., Thorup, K., & Tøttrup, A. P. (2019). Annual GPS tracking reveals unexpected wintering area in a long-distance migratory songbird. Journal of ornithology 160(1), 265–270.

Premuda, G., Gustin, M., Pandolfi, M., Sonet, L., & Cento, M. (2008). Spring raptor migration along the Adriatic coast (Italy): a comparative study over three sites. Avocetta 32, 13–20.

Rebassa, M., Machado, J., Martínez, J. L., Torrens, S. & Oriola, M. C. (2013). A Birding Tourist’s Guide to Mallorca. 2^nd^ Edition, 116–121.

Solt Sz. (2018). From arrival to mating. In: Palatitz, P., Solt, Sz., Fehérvári P. (Eds.) (2018). The Blue Vesper. Ecology and Conservation of the Red-Footed Falcon, Budapest, MME, 53–62.

Solt Sz. (2018). From egg-laying to hatching. In: Palatitz, P., Solt, Sz., Fehérvári P. (Eds.) (2018). The Blue Vesper. Ecology and Conservation of the Red-Footed Falcon, Budapest, MME, 63–72.

Somogyi, B.A., Horváth, G.F. (2019). Seasonal activity of common vole (*Microtus arvalis*) in alfalfa fields in southern Hungary. Biologia 74, 91–96

von Transehe N. & Schüz E. (1940) Massendurchzug des Rotfußfalken im Spätsommer 1939. Vogelzug (11), 31–35.

